# Increasing Rates of Diagnosis, Substantial Co-occurrence, and Variable Treatment Patterns of Eosinophilic Gastritis, Gastroenteritis and Colitis Based on 10 Year Data Across a Multi-Center Consortium

**DOI:** 10.1101/413583

**Authors:** Robert D. Pesek, Craig C. Reed, Amanda B. Muir, Patricia C. Fulkerson, Calies Menard-Katcher, Gary W. Falk, Jonathan Kuhl, Adam Z. Magier, Faria Ahmed, Maureen Demarshall, Ankur Gupta, Jonathan Gross, Tokunbo Ashorobi, Christina L. Carpenter, Jeffrey P. Krischer, Nirmala Gonsalves, Jonathan M. Spergel, Sandeep K. Gupta, Glenn T. Furuta, Marc E. Rothenberg, Evan S. Dellon, on behalf of the Consortium of Eosinophilic Gastrointestinal Disease Researchers (CEGIR)

## Abstract

**Financial Support and Acknowledgements:** Support for this project was provided through a research training grant as part of the Consortium of Eosinophilic Gastrointestinal Disease Researchers (CEGIR) (U54 AI117804). CEGIR is part of the Rare Disease Clinical Research Network (RDCRN), an initiative of the Office of Rare Diseases Research (ORDR), NCATS, and is funded through collaboration between NIAID, NIDDK, and NCATS. CEGIR is also supported by patient advocacy groups including APFED CURED and EFC. This project also received support from NIH T32 DK007634 (CCR).

**Author Disclosers:** Patricia Fulkerson: Grant funding from the NIH; Consultant for Genentech, Inc; Research support from Knopp Biosciences, LLC.

Gary Falk: Research support from Shire, Celgene, Adare, Regeneron. Consulting for Shire

Jonathan M. Spergel: Consultant for Regeneron, DBV Technology, Kaleo; Grant funding from DBV Technology, Aimmune Therapeutics, Food Allergy Research Education; Royalties from UpToDate

Nirmala Gonsalves: Royalties from UpToDate; Advisory board for Allakos

Sandeep K Gupta: Consultant for Alkalos, Abbott, QOL, Receptos; research support from Shire

Glenn Furuta: Founder of EnteroTrack; Consultant for Shire; Royalties from UpToDate

Marc E. Rothenberg: Consultant for Pulm One, Spoon Guru, ClostraBio, Celgene, Shire, Astra Zeneca, GlaxoSmithKline, Allakos, Adare, Regeneron and Novartis and has an equity interest in the first four listed and Immune Pharmaceuticals, and royalties from reslizumab (Teva Pharmaceuticals), PEESSv2 (Mapi Research Trust) and UpToDate. M.E.R. is an inventor of patents owned by Cincinnati Children’s.

Evan Dellon: Consultant for Adare, Allakos, Alivio, Banner, Celgen/Receptos, Enumeral, GSK, Regeneron, Shire; Research funding from Adare, Celegene/Receptos, Miraca, Meritage, Nutricia, Regeneron, Shire, Educational grant from Banner, Holoclara

**Study Highlights:** *What is current knowledge?:*

- Eosinophilic gastrointestinal disorders (EGIDs) include eosinophilic esophagitis (EoE), eosinophilic gastritis (EG), gastroenteritis (EGE), and colitis (EC).
- Non-EoE EGIDs are rare with most studies limited to case reports or review of single center experiences.
- There are no widely established guidelines for the diagnosis of EG, EGE, or EC.

*What is new here?:*

- In this multicenter study, EG, EGE, and EC were all diagnosed with increasing frequency over the past decade.
- Presenting symptoms are non-specific and do not reliably distinguish between disorders.
- There was no male predominance and the majority of subjects had atopy.
- Co-occurrence of EG, EGE, and EC diagnoses is common, seen in 41% of patients.
- There is substantial variability between centers in initial treatment approaches.

## Introduction

Eosinophilic gastrointestinal disorders (EGIDs) represent a group of diseases characterized by eosinophil-driven inflammation of specific locations in the GI tract^1-4^. The exact mechanisms that promote these disorders are not completely understood, but skewed Th2 immune response, frequently driven by food allergy, is believed to play a central role in recruiting eosinophils to the gastrointestinal tissue leading to dysfunction^5-6^. The best characterized EGID is eosinophilic esophagitis (EoE). Unlike the esophagus, eosinophils are a normal finding in other segments of the GI tract and are believed to play a role in the mucosal immune response. There can be variability in the number of tissue eosinophils depending on location. Elevations in gastrointestinal eosinophils can occur in a variety of settings such as drug or food allergies, parasitic infection, malignancy, inflammatory bowel disease, and hypereosinophilic syndrome^3-4^. If an increased number of eosinophils are found in the stomach, small intestine, or colon and other causes for the eosinophilia are ruled out, a patient is considered to have a non-esophageal EGID including eosinophilic gastritis (EG), enteritis (EE), gastroenteritis (EGE), or colitis (EC).

There are several factors that have hindered the understanding of non-esophageal EGIDs. Each is considered rare with an estimated prevalence from 2.1 to 8.2/100,000^7-8^. Current literature is predominately limited to case reports or single center retrospective studies with relatively small populations^9-16^. Also, there are also no published randomized, controlled treatment trials in these conditions. These factors limit the conclusions that can be drawn about etiology, risk factors, diagnosis, treatment response, and long-term outcomes. It has also been difficult to recognize these disorders as many of the presenting symptoms are protean, there is wide variability in the approach to endoscopic biopsies, and histologic findings are ill-defined^17^. Some patients may also have involvement of multiple segments of the gastrointestinal tract, but the significance of eosinophilic-predominant inflammation outside of the primary site of disease is unknown. As a result, additional studies are needed to better characterize non-esophageal EGIDs and address the current gaps in diagnosis and management which will enhance the ability for providers to care for these patients^18^.

The aims of this study were to determine the frequency of diagnosis, characterize the clinical features, assess the frequency of overlap in the conditions, and characterize initial provider management of non-esophageal EGID patients across a multi-site consortium over the past decade. We also hypothesized that the age at diagnosis would differ between non-esophageal EGIDs with EC having a younger age at diagnosis than EG or EGE and that sex distribution would be similar between disorders with no male predominance, which is seen in EoE.

## Methods

### Subject selection and areas of study

We conducted a retrospective study of subjects from multiple centers in the Consortium of Eosinophilic Gastrointestinal Disease Researchers (CEGIR)^19-20^, including pediatric and adult tertiary care centers in different geographical regions. Subjects were included from the University of Arkansas for Medical Sciences/Arkansas Children’s Hospital, the University of North Carolina, Children’s Hospital of Philadelphia, Cincinnati Children’s Hospital Medical Center, the University of Pennsylvania, and Children’s Hospital of Colorado. Each site reviewed clinical and research databases from 2005 to 2016 to identify patients with a non-esophageal EGID diagnosis. Subjects were considered for inclusion if they had a clinically confirmed diagnosis of EG, EGE, and/or EC by a provider from one of the study sites. Patients with EoE could be included if they had an additional EGID diagnosis. Eligible subjects were required to have clinical symptoms of gastrointestinal dysfunction and findings of increased eosinophils on GI tract biopsy at the time of diagnosis, before treatment was initiated. As there are no consensus diagnostic guidelines for non-esophageal EGIDs, when pathology reports enumerating eosinophil counts were available, these reports were reviewed to confirm the number of eosinophils reached a threshold level based upon the area of biopsy: stomach ≥ 30 eosinophils/high powered field (hpf); small intestine ≥ 50 eosinophils/hpf; or colon ≥ 60 eosinophils/hpf^21^. Not all centers had a report with the specific number of eosinophils and in these cases subjects were also considered eligible for inclusion if the pathologist’s description of the GI tract biopsy noted increased eosinophils with exclusion of other causes of gastrointestinal eosinophilia.

Data collected on subjects who met the inclusion criteria included demographics (age, gender, race, state of residence, and type of insurance), medical history of atopic and other co-existing diagnoses, symptoms at the time of diagnosis, and year of diagnosis (**Appendix 1**). The frequency of involvement of multiple segments of the GI tract was also determined. Data regarding treatments initiated at the time of diagnosis were also collected including use of medications and/or food elimination diets. These data were compared to determine center-to-center variability. All data were extracted from medical records and placed on a standardized spreadsheet. De-identified electronic data from each site was then submitted to a central data repository established by the CEGIR data management and coordinating center (DMCC) after which analysis was performed. This study was approved by NIH and NIAID, the CEGIR central Institutional Review Board (IRB) at Cincinnati Children’s Hospital Medical Center, as well as each of the participating site’s local IRBs.

### Statistical Analysis

For analysis, we first summarized the characteristics of the patient sample to examine the distribution of the variables and to assess for missing data and extreme values. The mean, standard deviation, and the shape of each distribution were calculated for continuous variables. Frequencies were tabulated for categorical variables. Bivariate analysis of categorical variables was conducted with Fisher’s exact test. Student’s t-test was used to compare continuous variables in two categories. One way analysis of variance was used to compare continuous variables with greater than two categories. All analyses were performed using Stata 14.2 (StataCorp, College Station, TX).

## Results

### Overall population and diagnostic time trends

A total of 376 subjects were enrolled including 317 children less than 18 years of age and 56 adults. The overall age range of patients was 0.5 to 77 years (**Table 1**). For children, the mean age was 7.3 years (range: 0.5-17) and for adults 35.9 years (range: 18-77). The overall population was 52% male and 71% of subjects were Caucasian. The second most common ethnicity was African American (10%). The majority of subjects had private compared to state/federal insurance (70% versus 18%, p = 0.02). The study population was highly atopic with approximately 60% of subjects having a history of at least one atopic condition (**Table 2**). In both children and adults, there was no significant differences between EGID diagnosis with regards to race, gender, or atopic history. Other common medical history included gastroesophageal reflux (22%), while conditions including celiac disease (3%), irritable bowel syndrome (2%), and inflammatory bowel disease (3%) were less common.

**Table 1.**
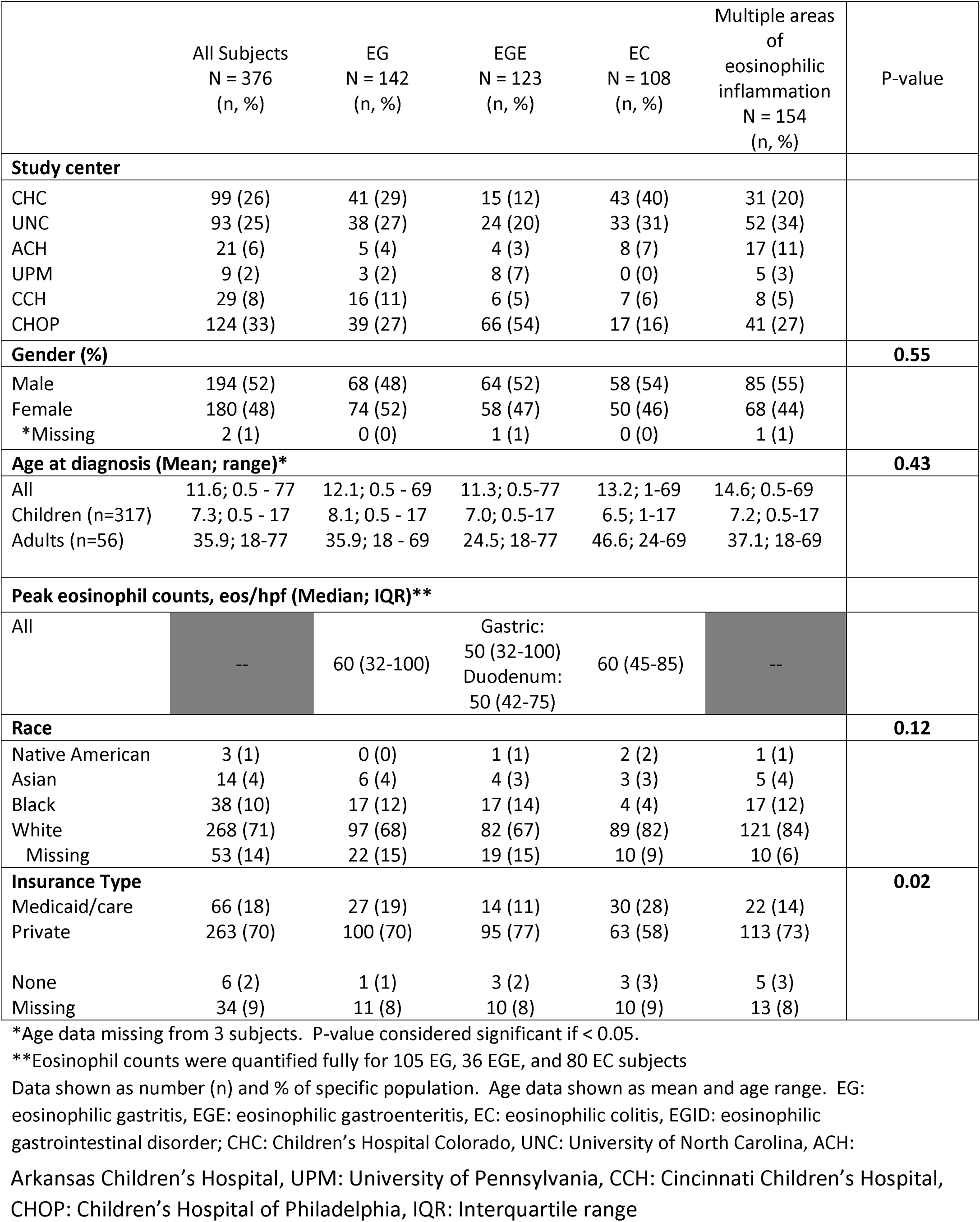
Demographics of study population and by EGID diagnosis (N = 376).

**Table 2.** Medical history of study population and by EGID diagnosis.

|  | All Subjects<br>N = 376<br>(n, %) | EG<br>N = 142<br>(n, %) | EGE<br>N = 123<br>(n, %) | EC<br>N = 108<br>(n, %) | Multiple areas<br>of eosinophilic<br>inflammation<br>N = 154<br>(n, %) |
| --- | --- | --- | --- | --- | --- |
| **Condition** |  |  |  |  |  |
| Any atopic condition | 221 (59) | 81 (57) | 90 (73) | 52 (48) | 96 (62) |
| Allergic conjunctivitis | 6 (2) | 3 (2) | 2 (2) | 2 (2) | 5 (3) |
| Allergic rhinitis | 87 (23) | 34 (24) | 38 (31) | 15 (14) | 48 (31) |
| Angioedema | 0 (0) | 0 (0) | 0 (0) | 0 (0) | 0 (0) |
| Asthma | 93 (25) | 38 (27) | 35 (28) | 24 (22) | 48 (31) |
| Atopic Dermatitis | 55 (15) | 18 (13) | 27 (22) | 11 (10) | 31 (20) |
| Drug Allergy | 42 (11) | 16 (11) | 15 (12) | 11 (10) | 12 (8) |
| Environmental allergy | 9 (2) | 3 (2) | 4 (3) | 2 (2) | 3 (33) |
| Food allergy | 117 (31) | 41 (29) | 56 (46) | 21 (19) | 54 (46) |
| Latex allergy | 4 (1) | 2 (1) | 2 (2) | 0 (0) | 3 (2) |
| Urticaria | 3 (1) | 2 (1) | 1 (1) | 0 (0) | 1 (1) |
| Venom allergy | 0 (0) | 0 (0) | 0 (0) | 0 (0) | 0 (0) |
| GERD | 82 (22) | 40 (28) | 24 (20) | 16 (15) | 32 (21) |
| Celiac disease | 11 (3) | 5 (4) | 2 (2) | 2 (2) | 3 (2) |
| Irritable bowel syndrome | 8 (2) | 4 (3) | 0 (0) | 2 (2) | 2 (1) |
| Ulcerative colitis | 6 (2) | 1 (1) | 0 (0) | 5 (5) | 1 (1) |
| Crohn's disease | 3 (1) | 1 (1) | 2 (2) | 1 (1) | 1 (1) |

Data shown as number (n) and % of specific population. EG: eosinophilic gastritis, EGE: eosinophilic gastroenteritis, EC: eosinophilic colitis, EGID: eosinophilic gastrointestinal disorder, GERD: gastroesophageal reflux disease.

When analyzing time trends, the frequency of diagnosis for all disorders increased throughout the study period, and this increase appeared to be most pronounced for subjects with multiple locations of eosinophilic inflammation outside of their primary disease site (**Figure 1**).

**Figure 1.**
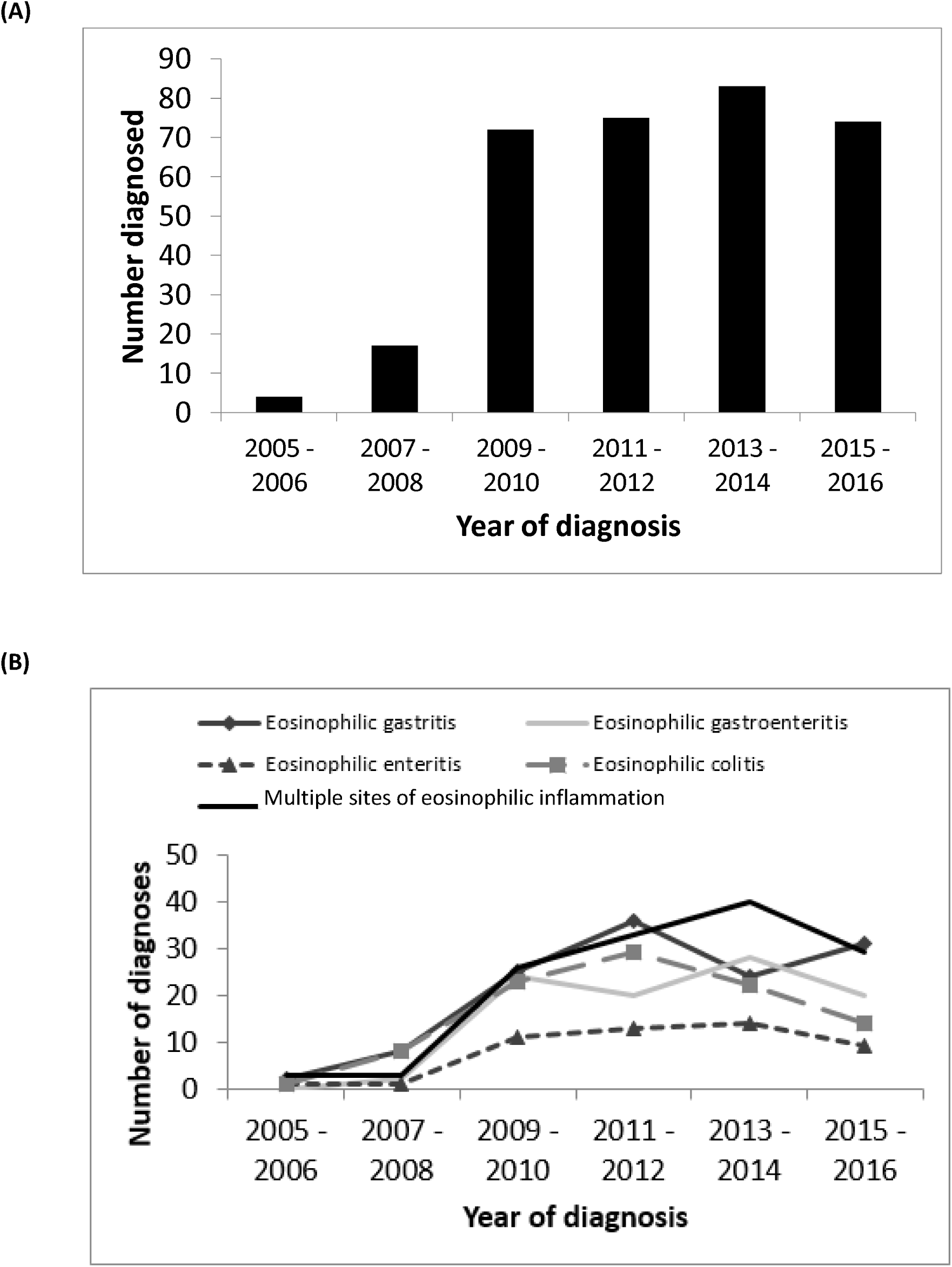
(A) Time trends for EGID diagnoses between 2005-2016 shorted by year and total number of diagnoses. (B) Frequency of EGID by diagnosis and year.

### EG, EGE, and EC

Each EGID was analyzed for frequency of diagnosis and clinical presentation. There were 142 subjects diagnosed with EG with slightly more females (52%) affected. Of these patients, 68% were Caucasian and 57% had a history of at least one atopic disease. The median peak gastric eosinophil count was 60 eos/hpf (n = 105; Interquartile Range (IQR): 32-100) (**Table 1**). The most common presenting symptoms included nausea/vomiting (54%) and abdominal pain (48%) (**Table 3**).

**Table 3.** Presenting symptoms by EGID diagnosis. Data shown as number (n) and % of specific population.

|  | All Subjects<br>N = 376<br>(n, %) | EG<br>N = 142<br>(n, %) | EGE<br>N = 123<br>(n, %) | EC<br>N = 108<br>(n, %) | Multiple areas<br>of eosinophilic<br>inflammation<br>N = 154<br>(n, %) |
| --- | --- | --- | --- | --- | --- |
| **Symptom(s)** |  |  |  |  |  |
| Headache | 7 (2) | 5 (4) | 3 (2) | 0 (0) | 3 (2) |
| Abdominal pain | 190 (51) | 68 (48) | 61 (50) | 65 (60) | 76 (52) |
| Regurgitation | 48 (13) | 26 (18) | 15 (12) | 6 (6) | 16 (10) |
| Irritability and/or crying | 18 (5) | 7 (5) | 2 (2) | 6 (6) | 4 (3) |
| Chest pain | 6 (2) | 4 (3) | 3 (2) | 2 (2) | 4 (3) |
| Nausea and/or vomiting | 185 (49) | 77 (54) | 64 (52) | 41 (38) | 87 (56) |
| Weight loss | 32 (9) | 13 (9) | 12 (10) | 8 (7) | 14 (9) |
| Eat too little or early satiety | 33 (9) | 18 (13) | 7 (6) | 9 (8) | 18 (12) |
| Nocturnal awakening due to pain | 3 (1) | 5 (4) | 1 (1) | 6 (6) | 2 (1) |
| Dysphagia | 42 (11) | 19 (13) | 9 (7) | 9 (8) | 29 (19) |
| Food impaction | 8 (2) | 3 (2) | 4 (3) | 0 (0) | 8 (5) |
| Food aversion/refusal | 31 (8) | 18 (13) | 5 (4) | 7 (6) | 10 (6) |
| Diarrhea | 113 (30) | 27 (19) | 39 (32) | 56 (52) | 42 (27) |
| Constipation | 58 (15) | 28 (20) | 14 (11) | 18 (17) | 20 (13) |
| Bloating | 10 (3) | 4 (3) | 2 (2) | 6 (6) | 7 (5) |
| Bloody stools | 40 (11) | 8 (6) | 7 (6) | 26 (24) | 14 (9) |
| No symptoms | 8 (2) | 2 (1) | 3 (2) | 1 (1) | 2 (1) |

EG: eosinophilic gastritis, EGE: eosinophilic gastroenteritis, EC: eosinophilic colitis, EGID: eosinophilic gastrointestinal disorder.

There were 123 subjects diagnosed with EGE, and of these, males (52%) and Caucasians (67%) were most affected. Atopy was also common with 73% reporting a history of at least one atopic disease. The median peak gastric eosinophil count for EGE subjects was 50 eos/hpf (n = 36; IQR: 32-100) and mean peak small intestine eosinophil count was 50 eos/hpf (n= 36; IQR: 42-75). Common presenting symptoms included nausea/vomiting (52%), abdominal pain (50%), and diarrhea (32%).

EC affected 108 subjects and was slightly more common in males (54%). As with the other EGIDs, Caucasians (82%) were most commonly affected. Subjects with EC also frequently noted atopic (48%) conditions. The mean peak colonic eosinophil count was 60 eos/hpf (n = 80; IQR: 45-85). Presenting symptoms were similar to the other diseases with abdominal pain (60%), diarrhea (52%), and nausea/vomiting (38%), however, a large portion of EC patients (24%) also presented with bloody stools.

In general, each EGID presented at a similar age in children (6.5 – 8.1 years)(**Table 1**). In adults, patients with EGE presented earlier (24.5 years) compared to EG or EC, while EC was typically diagnosed at later age (46.6 years) although this difference was not statistically significant. Across all EGID diagnoses, adults were more likely to present with dysphagia (p = 0.001) and diarrhea (p = 0.02) than children. Bloody stools were more common in subjects without an atopic history (p = 0.02) while females were more likely to have abdominal pain (p < 0.001).

### Multiple sites of eosinophilic inflammation

In the population studied, 154 subjects (41%) had additional locations of eosinophilic inflammation outside of their primary site of disease involvement (**Table 1**). Subjects with EGE were considered to have a single diagnosis and not two separate sites of inflammation. This was more common in children (n=116; 68%) than in adults (n=38; 37%) (p < 0.001). Multi-segment eosinophilic inflammation occurred in 55% of males and in 84% of Caucasians. Age at presentation was similar to single segment involvement (children: 7.2 years; adults: 37 years). Affected subjects were frequently atopic (62%) and presented similarly to those with isolated EG, EGE, or EC with nausea/vomiting (56%) and abdominal pain (52%) the most common symptoms (**Table 3**). Diarrhea (27%) and bloody stools (9%) occurred less frequently than in subjects with isolated EGE or EC. Of those with multiple sites of inflammation, 116 had esophageal involvement, and in these cases dysphagia was more common (23%) compared to inflammation limited to the organ of primary dysfunction. The most frequent combination of multi-site inflammation was the esophagus and stomach/small intestine (49 subjects) followed by the esophagus and stomach alone (42 subjects). Three subjects had involvement of the esophagus, stomach, and small/large intestines.

### Treatment Patterns

Initial treatment options were assessed for enrolled subjects with single segment disease (**Table 4**). For subjects with EG, the most commonly used medications included PPI (61%), topical corticosteroids (23%), and systemic corticosteroids (20%); 58% were treated with food elimination diets. In EGE, PPI (60%) were the most commonly prescribed medications followed by systemic corticosteroids (24%) and topical corticosteroids (20%); 68% were started on food elimination diets. For EC, PPI was the most commonly prescribed medication (30%), but 5-ASA (25%), systemic corticosteroids (19%), and enteric-coated budesonide (19%) were also frequently prescribed; 58% were treated with food elimination diets. When analyzing the types of food elimination diets utilized, the most commonly eliminated foods included milk (32%), soy (16%), egg (15%), and peanut (12%). Multiple concomitant treatments were used in 41%. Diet and corticosteroids were utilized in 27% of subjects with EG, 14% of subjects with EGE, and 31% of subjects with EC. Children and adults were treated with food elimination diets at similar rates (55% for each age group; p = NS) while adults were treated more frequently with combination diet and medication therapy. When comparing results by center, though PPI was the most common initial medication used for all centers and EGID diagnosis, there was a wide variability in approaches (**Figure 2**).

**Table 4.** Initial treatment of EGID by diagnosis sorted by type of medication and diet modification.

|  | EG<br>N = 142<br>(n, %) | EGE<br>N = 123<br>(n, %) | EC<br>N = 108<br>(n, %) | P-value |
| --- | --- | --- | --- | --- |
| **Medications** |  |  |  |  |
| PPI | 86 (61%) | 74 (60%) | 32 (30%) | < 0.001 |
| Topical corticosteroids | 33 (23%) | 24 (20%) | 11 (10%) | 0.008 |
| Systemic corticosteroids | 28 (20%) | 30 (24%) | 20 (19%) | 0.34 |
| Crushed enteric coated budesonide | 18 (13%) | 12 (10%) | 21 (19%) | 0.016 |
| 5-ASA | 1 (1%) | 4 (3%) | 27 (25%) | < 0.001 |
| Immune modulating drug | 5 (4%) | 4 (3%) | 8 (7%) | 0.34 |
| Mast cell agent | 1 (1%) | 2 (2%) | 4 (4%) | 0.65 |
| Monoclonal antibody | 2 (1%) | 4 (3%) | 1 (1%) | 0.15 |
| **Diet Modification** |  |  |  |  |
| Six-food elimination | 5 (4%) | 2 (2%) | 6 (6%) | 0.43 |
| Allergen specific elimination | 62 (44%) | 62 (50%) | 43 (40%) | 0.31 |
| Elemental diet | 15 (11%) | 20 (16%) | 12 (11%) | 0.17 |

Six food elimination included restriction of cow's milk, hen's egg, soy, wheat, nuts, fish, and shellfish. Data shown as number (n) and % of specific population. P-value considered significant if < 0.05. EG: eosinophilic gastritis, EGE: eosinophilic gastroenteritis, EC: eosinophilic colitis, EGID: eosinophilic gastrointestinal disorder, PPI: proton-pump inhibitor, 5-ASA: 5-aminosalicylic acid.

**Figure 2.**
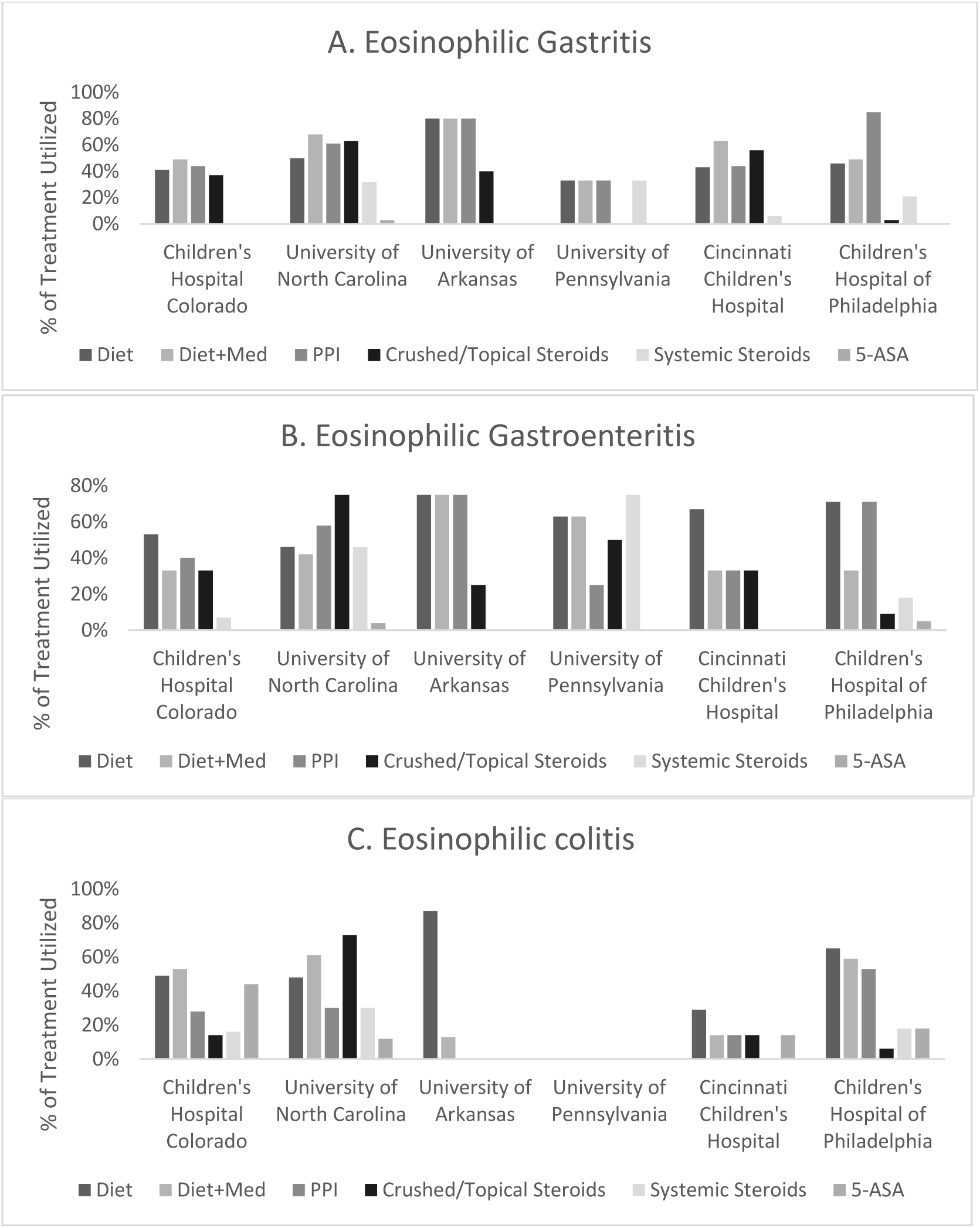
Initial medical treatments sorted by center and EGID diagnosis demonstrating high variability in treatment approaches. Bars represent percentage of treatment utilized; subjects could receive more than one treatment per disease. (A) Initial treatments utilized for eosinophilic gastritis. PPI was the most commonly used medication (61%) followed by topical steroids (23%) and systemic steroids (20%). 58% were treated with food elimination diets. (B) Initial treatments utilized for eosinophilic gastroenteritis. PPI was the most commonly used medication (60%) followed by systemic steroids (24%). 68% were treated with food elimination diets. (C) Initial treatments utilized for eosinophilic colitis. PPI was the most commonly utilized medication (30%) followed by 5-ASA (25%), systemic steroids (19%), and enteric-coated budesonide (19%). Food elimination diets were utilized in 58% of cases. Multiple treatments were initiated in 41% of subjects across all disorders. PPI: proton-pump inhibitor, 5-ASA: 5-aminosalicyclic acid.

## Discussion

Because the non-esophageal EGIDs are rare disorders, data regarding these conditions are limited and even less is known about patients affected in multiple GI segments. This study examines the largest populations of non-esophageal EGIDs to date and there are several notable findings. First, each of these disorders has increased in frequency over the past decade at the participating centers. Second, over 40% of our population had eosinophilic inflammation of more than one GI segment. Inflammation affecting the entire intestinal tract occurred in approximately 1%. Third, unlike EoE, non-esophageal EGIDs do not have a strong male predominance. However, similar to EoE, Caucasians were most commonly affected. Fourth, there was a high level of concomitant atopy in all EGID subtypes, suggesting that these conditions, like EoE, may be allergen-mediated in many cases. Fifth, the clinical presentation is similar across each disease with nausea/vomiting and abdominal pain representing the most common symptoms. Patients with EC are frequently present with diarrhea and bloody stools which may help in recognition of this disorder. Finally, combination treatment for initial management of these disorders was common, with frequent use of both medications and food elimination diets. Significant center-to-center variability and lack of any single predominant treatment likely speaks to the heterogeneity of disease presentation and lack of data supporting the most effective treatment for each of these disorders.

A primary finding of the study is that the overall frequency of non-esophageal EGIDs is increasing. While two studies have used large insurance medical and pharmaceutical claims databases to estimate the prevalence of these diseases to be between 2.1 and 8.2/100,000, they were not able to assess trends of new diagnoses over time^7-8^. These studies also had an adult predominance in their database, and our population differs in that it was predominately pediatric subjects, which reflects the patient population at the participating sites. Neither incidence nor prevalence could be determined in our study as we were looking at individual cases from referral centers, and it is therefore difficult for us to draw conclusions on the relative frequency of EG, EGE, or EC in children compared to adults.

A second finding isthe presence of eosinophilic inflammation in locations other than the primary site of disease; this is of note as there are very few reports describing this phenomenon and none in a large cohort of subjects. The one exception is EGE. By definition, this represents an overlap of eosinophilic inflammation in both the stomach and small intestine. Although disease can be isolated to either organ, they are frequently seen together and in many reports are described together as a distinct diagnosis. The frequency of overlap involving other segments is less described. In a study by Reed et al. of 44 EGID patients, 30% of EGE patients had esophageal involvement, and 28% had colon involvement^22^. Choi et al. evaluated 24 children with EGE and noted concomitant esophageal involvement in 13%, colon involvement in 29%, and involvement of multiple segments in 54%^23^. Caldwell et al. found that 87% of EG patients had eosinophilia at other gastrointestinal sites^24^. Several other studies have found rates of eosinophilic inflammation at multiple sites varying from 20-88%^25-28^. Despite these reports, the overall prevalence of this phenomenon is unknown. This finding suggests that diagnostic workup should potentially encompass the entire GI tract, even in patients presenting only with upper GI symptoms. The presence of diarrhea and/or bloody stools may help identify patients who need colonoscopy, but in the study population, a large number of EC cases presented without these clinical symptoms. It is also unclear if inflammation affecting more than one segment of the GI tract represent a different disease state. In the study by Caldwell and colleagues, a distinct genetic signature was found for EG as compared to EoE, but similar data are not yet published on EGE or EC^24^. This type of study also has not been performed in subjects with multiple sites of inflammation. As each EGID represents a distinct disease, different diagnostic approaches may be needed and patients may respond differently to treatment.

In this study, patients with non-esophageal EGIDs were predominantly Caucasian with a high rate of atopy. These findings are supported by several other studies in both EoE and non-esophageal EGIDs^29-31^. In a study by Guajardo et al. utilizing a world-wide-web registry to evaluate demographics, presenting symptoms, and medical history, 80% of included subjects had atopy, including those with non-esophageal EGIDs^25^. Interestingly, multi-segment inflammation was also more common in patients with an atopic history. When compared with the results by Caldwell et al. in their study of EG patients, it supports their finding that non-esophageal EGIDs are Th2 driven diseases^24^.

We did not find a male predominance typically seen in studies of isolated EoE. Our findings are similar to others performed in non-esophageal EGID populations. In the study by Jensen et al., there was a female predominance in all non-esophageal EGIDs, but most pronounced for EG^7^. In a similar study by Mansoor et al., a similar female predominance was found in both EGE and EC^8^. Some studies have demonstrated a male predominance of certain non-esophageal EGIDs such as EC, but in general there appears to be a lack of male predominance in these disorders^32-33^.

When evaluating the initial management of these disorders, wide variability existed amongst participating centers, which may speak to differences in provider preference/experience, the lack of high level evidence to support any given therapy, as well as to the heterogeneity of the presentation of these disorders. PPIs were the most commonly used medication, even in patients with colonic disease. Given that the majority of patients presented with nausea/vomiting or abdominal pain, it is not surprising to see such a high rate of PPI use. Several different formulations of corticosteroids were also frequently used depending on the GI segment involved. Food elimination diets were common, which may reflect the large pediatric population included. Combination treatments, utilizing multiple medications or medication and diet were the most frequent initial approach across all sites. Clearly, larger trials are needed to determine the most effective treatment for non-esophageal EGIDs.

There are several limitations to acknowledge. First, this was a retrospective study with the inherent limitations of this design. Importantly, this prevented us from prospectively obtaining and reviewing a comprehensive set of biopsy samples. To counter this, the study design allowed for inclusion of subjects if the site pathologist reported an increased number of eosinophils on GI tract biopsy that was above normal thresholds. If the eosinophil count was not specified, the subject was enrolled only if there was a confirmed clinical diagnosis. It is still possible that subjects may have been included that do not meet the currently accepted criteria for diagnosis of EG, EGE, and/or EC. For instance, the number of fields used to specify tissue eosinophil counts was not evaluated and sites may not have used peak eosinophil counts in 5 different tissue fields as recommended for EG^21^. Study inclusion was based upon a combination of provider diagnoses of EGID, associated clinical symptoms, and pathologic findings which should have reduced the risk of subjects enrolling who had another explanation for the findings of intestinal eosinophilia; this overall approach mirrors the way non-esophageal EGIDs are currently clinically diagnosed. Because data were obtained from referral centers, there could also be bias in regards to patient selection, but having the patients seen, diagnosed, and treated at expert centers lends confidence to the correct diagnosis. Finally, this was a pediatric predominant population with much less adult representation based on the participating sites. However, when comparing adults versus children, we did not find significant differences in gender, race, medical history, or presenting symptoms.

In conclusion, in this relatively large multicenter retrospective study, diagnoses of EG, EGE, and EC are increasing. Non-esophageal EGID subjects are frequently Caucasian with a history of atopy, however, there is no male predominance as seen in EoE. Further, many subjects are affected by inflammation in segments of their GI tract outside of the site of primary disease. This may be one explanation for why the clinical presentation is similar between the disorders. More work is needed to understand the significance of multi-segment inflammation and to determine if this represents a distinct clinical entity. Further, the atopic predisposition suggests that these conditions, like EoE, may well be allergen- or immune-mediated, and future work should address this hypothesis. There is also significant variability in regards to treatment of these disorders. Larger and prospective studies are need to determine effective diagnostic algorithms and treatment strategies.

